# From Growth Faltering to Recovery: Gut Microbial and Body Composition Signatures of Early Childhood Malnutrition Phenotypes

**DOI:** 10.64898/2026.05.11.723332

**Authors:** Evelyn Takyi, Douglas Momberg, Rihlat Said Mohamed, Jonathan Y Bernard, Shane A Norris, Linda Richter, Julian May, Alexia J Murphy-Alford, Lutricia Rakgoale, Venesa Sahibdeen, Claudine K. Nkera-Gutabara, Ovokeraye Oduaran, Rosa Krajmalnik-Brown, Lee E. Voth-Gaeddert

## Abstract

**Background:** Chronic malnutrition in early childhood is a multifactorial condition associated with long-term impairments, yet the physiological and gut microbial pathways underlying differential growth trajectories remain poorly understood.

**Objective:** We aimed to characterize phenotypic growth trajectories and identify the associated gut microbial and body composition signatures in infants during the first year of life.

**Methods:** We analyzed longitudinal data from birth to 12 months in a South African cohort (Soweto, n=45). Individual linear growth trajectories were modeled using the Jenss-Bayley equation, and children were clustered based on model parameters to identify phenotypic subgroups. Body composition (fat-free mass and fat mass) was measured via deuterium dilution at 6 and 12 months, and gut microbiome development was assessed using 16S rRNA gene amplicons at 4, 6, and 12 months.

**Results:** We identified distinct phenotypic subgroups including healthy growth, catch-up growth, and growth faltering, that were obscured at the cohort level. These trajectories diverged most dynamically within the first 6 months of life. Integrated analysis revealed that in the growth faltering cluster, height-for-age and fat-free mass z-scores stabilized between 6 and 12 months, whereas fat mass z-scores (FMZ) declined. This trade-off is consistent with a catabolic state where energy reserves are prioritized for lean tissue and bone growth. Furthermore, at 6 months, the growth faltering cluster was enriched with opportunistic pathobionts (e.g., *Paraclostridium*). In contrast, the catch-up cluster exhibited a transient enrichment of facultative anaerobes (e.g., *Enterobacter*), supporting a hypothesis that these oxygen-tolerant taxa may help bridge a transitional microbial state in partially oxygenated or inflamed environments to enable physiological recovery.

**Conclusions:** Early childhood chronic malnutrition phenotypes in South African infants can be defined by distinct microbial and body composition signatures that diverge within six months of life. Integrated interventions should target both host anabolic state and microbiome transitions to support recovery.

## INTRODUCTION

Chronic malnutrition remains a pressing global health challenge, affecting 23.2% of children globally in 2024 (Black et al., 2013; UNICEF/WHO/World Bank Group, 2021; United Nations Children’s Fund (UNICEF) et al., 2025; Victora et al., 2021). Chronic malnutrition, or stunting, is defined as being two or more standard deviations below the World Health Organization (WHO) Child Growth Standard median for height-for-age (HAZ) (Members of the WHO Multicentre Growth Reference Study Group, 2006). Chronic malnutrition arises from a network of underlying factors, including inadequate dietary intake and environmental exposures (Black et al., 2013; Momberg et al., 2021; United Nations Children’s Fund (UNICEF) et al., 2025; Victora et al., 2021; Voth-Gaeddert et al., 2018). It is associated with short and long-term health challenges including morbidity, development (e.g., growth, neurodevelopment, metabolic), and increased risk for cardiometabolic diseases (Prendergast & Humphrey, 2014; Victora et al., 2008). Reducing the prevalence of childhood stunting remains a major global health priority (Vaivada et al., 2020). The determinants of stunting and other forms of child undernutrition are complex. Recent nutritional and exposure-reduction interventions remain limited in their effect on child growth (Dewey & Begum, 2011; Pickering et al., 2019; Prendergast & Humphrey, 2014). A study conducted in India by Taneja et al. showed that health, nutrition, psychological care and improved water, sanitation and hygiene (WASH) interventions delivered during preconception and pregnancy periods reduced the low birth weight and stunting at 24 months (Taneja et al., 2022). However, a study by Humphrey et al. in Zimbabwe showed that improved WASH and Infant and young child feeding (IYCF) interventions may not reduce chronic malnutrition (Humphrey et al., 2019). Another study conducted by Null et al. and Luby et al. showed that WASH had no effect on linear growth of children aged 1 to 2 years born in Kenya (Null et al., 2018) and Bangladesh (Luby et al., 2018). This suggests that other factors, such as the gut microbiome and body composition, may be important components in understanding the key mechanisms that drive proper growth (Blanton et al., 2016; Lumley et al., 2026).

Emerging evidence supports the role of the commensal gut microorganisms in mediating child growth (Cheng et al., 2023; Chibuye et al., 2024; Voth-Gaeddert et al., 2019). Microbes influence a number of metabolic, immune, and endocrine pathways in early life that contribute to early-life growth and development (Nirmalkar et al., 2022; Robertson et al., 2019) suggesting that disturbances to common microbiome maturation characteristics may impair critical growth and developmental pathways (Subramanian et al., 2014). Furthermore, body composition, fat mass (FM) and fat-free mass (FFM), also have unique roles in early childhood growth (Momberg et al., 2023; Wells, 2019). Recent studies suggest that FFM accretion is more strongly associated with linear growth, while FM plays a more variable role depending on age, infection burden, and nutritional context (Admassu et al., 2017). Emerging evidence also indicates that disruptions in early FFM development may predict persistent growth deficits, even in children who experience partial catch-up growth later in infancy (Fenton et al., 2024). However, limited data exists on integrating longitudinal data on growth, body composition, and the gut microbiome to gain a fuller understanding of child growth. Data suggests that dynamics of growth faltering and catch-up growth may occur at certain ages, under certain conditions, but little is known about the underlying physiological conditions contributing to unique subgroups that do or do not experience these dynamics.

In this study we leveraged longitudinal data collected from children born in Soweto, South Africa that include growth, body composition, gut microbiome data, morbidity, and environmental exposures among a cohort of children from birth to 12 months. This work aims to 1) identify unique growth phenotypes based on growth trajectories among children from birth to 12 months and 2) characterize differences in body composition, gut microbiome, and morbidity according to growth trajectory patterns.

## MATERIALS AND METHODS

### Study design

This study used data collected from a prospective cohort study in Soweto, South Africa, described previously (Momberg et al., 2020). Briefly, it consisted of mother-infant pairs (n=45), *a priori* born healthy, that were recruited and followed up weekly from birth to 6 months and fortnightly from 6 to 12 months postnatally. Recruitment commenced in January 2018, and data collection ended when the last recruited infant turned one year of age in March 2019. All the participants were screened and recruited at the maternity services at Chris Hani Baragwanath Academic Hospital in Soweto, Johannesburg. The inclusion criteria were women who were 1-3 days postpartum, 18 years and above of age at the time of screening, singleton pregnancy, birthweight between ≥2,500 and <4,000 g, and term pregnancy between ≥38 and <42 weeks gestational age. The exclusion criteria were mother-infant pairs diagnosed with physical, mental, or congenital disorders at birth or if a mother was living with HIV. Other study settings and design details are reported in Momberg et al. (Momberg et al., 2020). The study protocol was approved by the University of the Witwatersrand’s Human Research Ethics Committee (Medical) (Certificates: M170753, M170872, and M170955).

### Exposure and Outcome variables measurements

A random subset of participants in the cohort study were selected for additional data collection (n=45 mother-infant pairs). Briefly, anthropometric data (sex, age, length, weight) related to infants’ growth were collected following WHO standardized protocols (Members of the WHO Multicentre Growth Reference Study Group, 2006) and z-scores generated via the WHO Anthro Survey Analyser software (World Health Organization, 2020). Infants’ body composition data (FM and FFM) was collected using the International Atomic Energy Agency (IAEA) deuterium oxide dilution technique described previously (International Atomic Energy Agency, 2010; Sakamornchai et al., 2023). Trained professionals orally administered 1 mL of deuterium oxide to the infants, and saliva samples were collected as follows: 2 mL prior to deuterium dose (baseline), 2 mL sample at 2h:30min post-dose, and a final 2 mL sample at 3 hours post-dose. This was conducted at 6 and 12 months of age. Results were analyzed using an Agilent 4500 Series spectrometer. FM and FFM were calculated from total body weight (TBW) and were adjusted for height and presented as FM Index (FMI) and FFM index (FFMI). FM and FFM Z scores (FMZ and FFMZ) were generated from the IAEA MIBCRS Body composition reference charts (Murphy-Alford et al., 2023).

Exposure variables relating to infant morbidity and illness data were collected using an interviewer-administered questionnaire and confirmed against clinical records at six months. Household water, sanitation, and hygiene (WASH) was assessed using a questionnaire adapted from the WHO/UNICEF(WHO & UNICEF, 2006; WHO/UNICEF Joint Monitoring Programme (JMP), 2015; World Health Organization, 2019). The details of the questionnaires collected on water, sanitation, and hygiene indices are documented and reported in (Momberg et al., 2020). Other variables relating to maternal anthropometry and infant feeding practices were also reported in Momberg et al. (Momberg et al., 2020).

### Analysis of growth data of infants using Jenss Bayley Model

The Jenss-Bayley growth model was used to assess individual growth trajectories using height data of infants from birth to 12 months, collected weekly from 0 to 6 months and fortnightly from 6 to 12 months (Jenss & Bayley, 1937). The Jenss-Bayley model consists of fitting a four parameter mixed-effect model to physical growth data to obtain individual growth curves and parameters (Berkey, 1982). Briefly, the Jenss-Bayley model (y = a + bt - e^c+dt^) utilizes four interpretable parameters to capture growth from birth through age 8. Parameter A serves as the intercept (representing the measure at birth), while Parameter B represents the stable growth velocity established after the initial infancy period. The early childhood growth spurt is captured by Parameter C, reflecting the magnitude of the infant spurt, and Parameter D, which represents the rate of growth deceleration as the child transitions toward a linear trajectory. The output of the Jenss Bayley model for each of the parameters was used to conduct a k-means cluster analysis and visualized via Principal Component Analysis to identify infants with similar features of growth patterns. The k-means cluster analysis was conducted on the four estimated parameters while the Elbow and Silhouette methods were used to identify the optimal number of clusters. This was conducted in R and details can be found in the supplemental material (Fenton et al., 2024).

### Microbiome Study

#### Sample collection and 16S rRNA gene amplicon sequencing

Details on the sample collection procedure for microbiome analysis were described in detail in a dissertation by Nkera-Gutabara (Nkera-Gutabara, 2020). Briefly, child stool samples were collected at three time points (4, 6, and 12 months) for each of the 45 participants using the OMNIgene gut collection tubes (DNA Genotek, 2017). The tubes used enabled the accessible collection and contained a buffer for the stabilization of stool samples for gut microbiome profiling by rapidly homogenizing and stabilizing the sample at the time of collection. A total of 112 fecal samples were collected. DNA extraction from the child stool samples was performed using the QIAamp Power Faecal DNA kit with slight modifications to the protocol to increase the yield. Amplification of the microbial 16S rRNA gene was carried out at the National Institute for Communicable Diseases (NICD) Sandringham, Johannesburg sequencing core facility. PCR amplification of the V3 and V4 genomic regions of the 16S rRNA gene was performed and sequenced on an Illumina MiSeq (Illumina Inc., San Diego, California, USA) according to standard manufacturer protocols. Demultiplexed read pairs from the MiSeq runs were analyzed using the Quantitative Insights Into Microbial Ecology 2 (QIIME2) software version 2019.7 (Bolyen et al., 2019). The DADA2 pipeline was used to process Amplicon Sequence Variants (ASVs) (Callahan et al., 2016). ASVs were classified to the lowest possible taxonomic rank using QIIME2 and reference dataset from the SILVA database v.132 (Quast et al., 2013).

#### Bioinformatic Analysis

The statistical software used to analyze all the data was R version 4.0.2 (R Core Team, 2020). The taxonomic composition data were rarefied to account for differences in sequencing depth of 20469 reads per sample. Alpha-diversity metrics and beta-diversity (Bray–Curtis dissimilarity index) were calculated to explore taxonomic diversities. Univariate analysis was performed for alpha diversity and visualized using the ggpubr R package (v0.4.0). For beta diversity, permutational multivariate analysis of variance (PERMANOVA) with the “adonis” function and Bray-Curtis dissimilarity was calculated with the “vegan” package and used for principal-coordinates analysis (Oksanen et al., 2001). All p-values were corrected for multiple hypothesis comparisons with the Benjamini–Hochberg method. The differential abundant taxa were evaluated using the linear discriminant (LDA) effect size (LEfSe) analysis method to determine taxa that significantly differed in abundance (Segata et al., 2011) as previously done (Zeibich et al., 2021).

## RESULTS

### Analysis of the HAZ and Body Composition in the infant’s population revealed four subgroups (clusters)

The overall distributions for all children for HAZ and weight-for-height z-scores (WHZ) are depicted in Figure S1 and S2. There were no significant differences between measurement timepoints at birth (0 months), 6 months, or 12 months. However, using k-means clustering applied to child-specific parameters from the Jenss Bayley equation, we identified four subgroups (Figure 1A). Factor loadings from the PCA suggested that PC1 primarily loaded on parameters B (late growth dynamics), C (inflection point in growth curve) and D (early rapid growth) while PC2 primarily loaded on parameter A (initial height). Figure 1B depicts the HAZ scores, FM and FFM at birth, 6, and 12 months for each cluster. Statistically significant differences were observed in HAZ scores between the four clusters (Supplementary Figure S2A) and between timepoints within clusters 3 and 4 (see Figure 1B). The data suggest that there were unique subgroup growth trajectories not apparent when analyzing all aggregated data.

**Figure 1.**
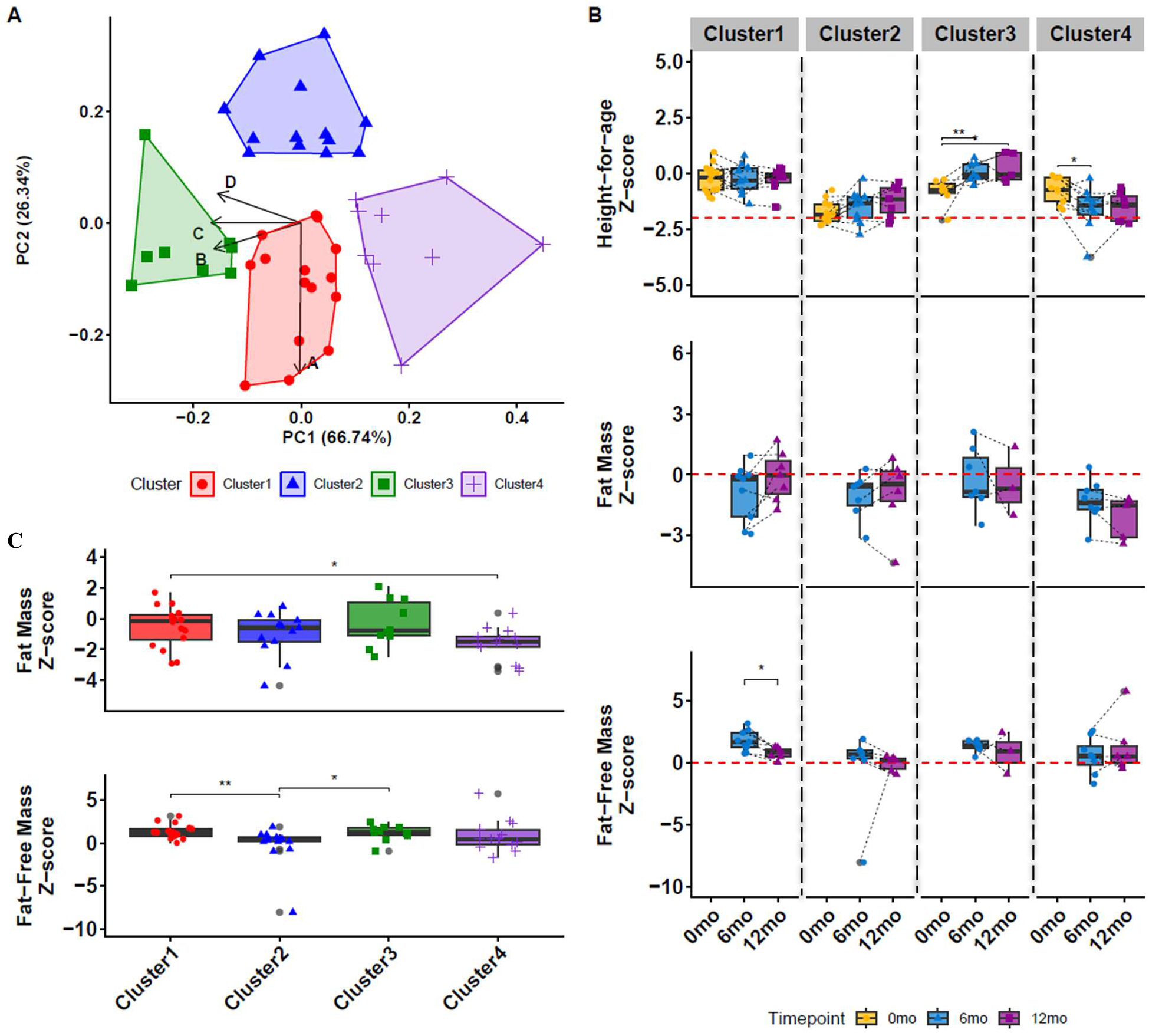
**A)** Principal Components Analysis (PCA) of children clustered by the Jenss-Bayley parameter estimates. Factor loadings indicate PC1 represents B (late growth dynamics), C (inflection point in growth curve) and D (early rapid growth) while PC2 presents parameter A(initial height). **B)** Box plots of HAZ scores and body composition z-score metrics between timepoints within each identified cluster. **C)** Box plots of body composition z-score metrics within each identified cluster. ** p value < 0.01; * p value < 0.05; Wilcoxon rank sum tests were calculated, and all p-values were corrected with Benjamini-Hochberg post-hoc corrections.

Based on these cluster-specific growth trajectories, we attributed general descriptive ‘phenotypic’ labels to each. Individuals in cluster 1 maintained their HAZ scores over time and were classified as ‘healthy’. Individuals in cluster 2 started with a low HAZ score (∼-2) but slowly recovered over time at each timepoint (though not significant) and were classified as ‘stunted but recovering’. In cluster 3, individuals had lower HAZ scores at birth (∼-1) but recovered completely by 6 months of age and maintained this recovery through 12 months. Cluster 3 was classified as ‘catch up growth’. Finally, in cluster 4, infants were born with a lower HAZ score, similar to cluster 3, but worsened by 6 months, but then maintained this status through 12 months. Cluster 4 was classified as ‘growth faltering’. These trends are similar for the weight-for-age z-scores (WAZ) (underweight) (Supplementary Figure S2D) but varied slightly for WHZ (acute malnutrition; Figure S2E).

In the overall study population, there were no significant differences in the fat mass index (FMI) or fat mass z-score (FMZ-score) between 6 and 12 months (Supplementary Figure S3A, S3B, S3C) with 19% and 21% of children having an FMZ-score less than −2 at 6 and 12 months, respectively. For FFMI and FFMZ, there was a significant decrease between 6 and 12 months (Supplementary Figure S3D, S3E, S3F), however, only one child had a FFMZ less than −2, occurring at the 6-month time point. Next, comparing subgroups, there was a significant difference in overall FMZ between cluster 1 (healthy) and cluster 4 (growth faltering) (Figure 1C). When disaggregating by 6- and 12-month timepoints for FMZ, clusters 1, 2, and 3 trend upwards or neutral while cluster 4 trends downwards though not significant (Figure 1B). For FFMZ disaggregated by timepoint, clusters 1, 2, and 3 trend downward while cluster 4 appears neutral (Figure 1B). Finally, comparing overall body composition metrics to anthropometrics there was a significant correlation between FM (all metrics) and WAZ and WHZ at 6 and 12 months. However, there was no correlation between FFM (all metrics) and HAZ at either 6 or 12 months (see Supplementary Figure S5).

### Gut microbiome diversity was different between subgroups

The overall gut microbiome alpha and beta diversity were evaluated at the study population level and for each subgroup of children (see Figure 2 and Supplementary Figure S6). For the subgroup analysis by time point, there was a significant decrease in Shannon diversity (species richness + evenness) in clusters 3 and 4 between 4 and 6 months while all groups had a slight, non-significant increase between 6 and 12 months (Figure 2A). For richness (i.e., number of unique taxa), most children had a high richness score by 4 months in comparison to other timepoints (Figure 2B). Finally, we observed a decrease in evenness for all groups between 4 and 6 months, with at least partial recovery by 12 months (Figure 2C). Interestingly, cluster 4 had the largest changes in evenness between timepoints. At the study population level, there were no significant correlations between alpha diversity and any anthropometrics (see Supplementary Figure S7). For beta diversity, visual inspection of the Principal Coordinates Analysis (PCOA) plot suggests minimal separation for the 4 subgroups of children (Figure 2D). However, PERMANOVA analysis identified clusters that were significantly different in their microbial community among each other (Supplementary Table S6). Of the cluster comparisons that significantly differed in their microbial community, cluster 4 was significantly different from all other clusters while cluster 2 and 3 were significantly different from each other.

**Figure 2.**
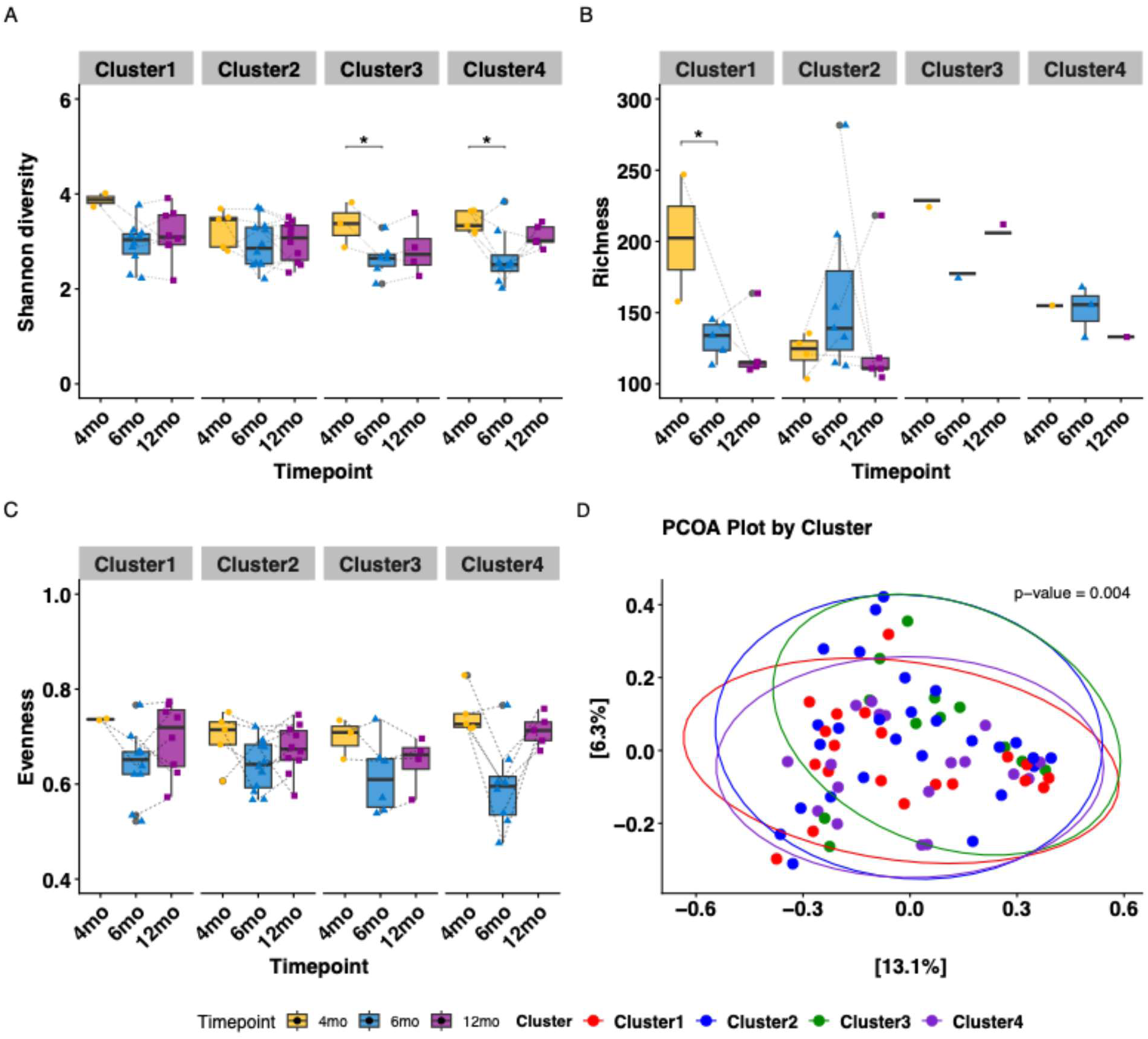
Gut microbiome diversity (alpha and beta diversity) plots by cluster and timepoints. A) Shannon diversity B) Richness C) Evenness D) Principal coordinates Analysis (PCOA) plot by Cluster based on a Bray-Curtis dissimilarity matrix. Wilcoxon signed-rank test was used to analyze the difference in alpha diversity measures. All p-values were corrected with Benjamini-Hochberg post-hoc corrections. * p<0.05 . P-value on PCOA plot indicate the significance of grouping with adonis2 Permutational Multivariate Analysis of Variance Using Distance Matrices test (PERMANOVA).

Next, we compared taxa-level differences across clusters, grouping all timepoints to understand what microbial communities might be consistently unique across all timepoints to 1) children experiencing growth faltering (Cluster 4), 2) children going through catch-up growth (Cluster 3 and Cluster 2), and 3) children who maintain good health (Cluster 1). For Cluster 4, we identified that sequences of *Paraclostridium* were more abundant compared to the other three clusters, while sequences of *Megasphaera* were potentially depleted compared to cluster 1 and cluster 2 (Figure 3A). This was supported further by evaluating the individual relative abundances of the sequences of these two taxa (Figure 3B).

**Figure 3.**
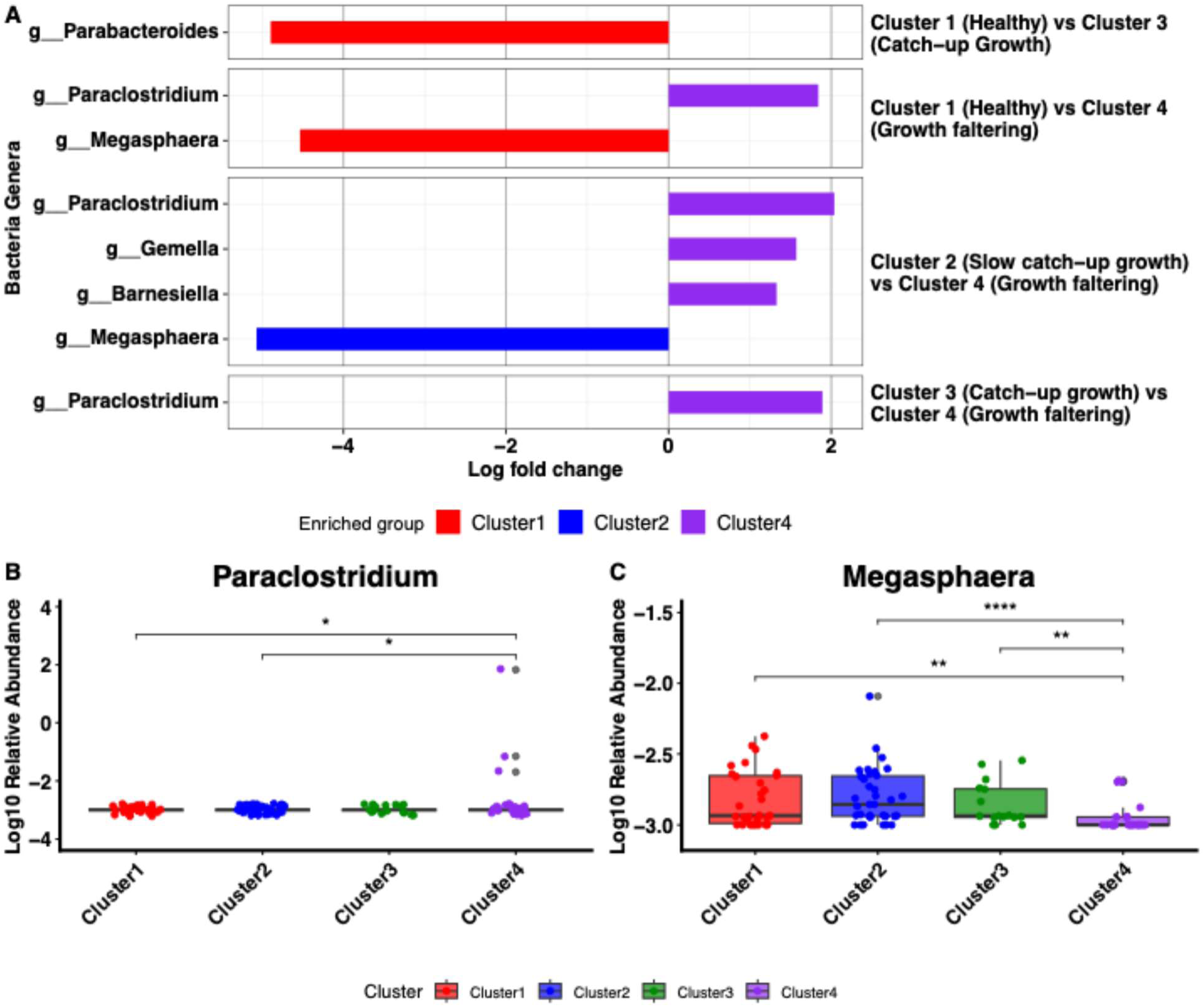
Comparison of relative abundance of bacterial Taxa. **A)** Differentially abundant taxa between cluster groups, assessed using LEfSe. The x-axis shows the log fold change representing the degree of differences in the relative abundance of the taxa between the cluster groups expressed in logarithmic scale. Only taxa having a p<0.05, corrected for multiple hypothesis with Benjamini-Hochberg post-hoc corrections and log fold change >1 are shown. The y-axis shows differentially abundant taxa between the cluster groups and is colored based on the cluster group in which they are most enriched. **B) and C)** Relative abundance (after log10 transformation) of *Paraclostridium* and *Megasphaera*, respectively.

### Unique gut microbes were associated with catch-up growth and growth faltering at 6 months

Given the dynamic nature of the gut microbiome in children, we also evaluated differences of the gut microbiome of these clusters at each time point as well (4, 6, and 12 months). Our goal was to better characterize unique taxa for the key growth phenotypes represented by the different clusters, with a focus on Cluster 1 (healthy), 3 (catchup growth), and 4 (growth faltering). While cluster 2 demonstrated some catchup growth, it was non-significant and accrued at a slower rate therefore we focused only on cluster 3 for catchup growth. While there were no significant differences in beta diversity between clusters at any individual time points, several differentially abundant taxon were identified (Supplementary Table S7 and Figure S8).

Given the early dynamic nature of growth identified in Figure 1, the 4-month and 6-month timepoints are of highest interest. At 4 months, the only differentially abundant taxon were *Actinomyces,* which had a higher abundance in Cluster 3 (catchup growth) compared to Cluster 4 (growth faltering). At 6 months, Cluster 1 (healthy) had a higher abundance of *Parabacteroides* compared to Cluster 3 (catchup growth) and Cluster 4 (growth faltering). For Cluster 3 (catchup Growth), *Streptococcus*, *Klebsiella*, and *Enterobacter* were higher in abundance compared to both Cluster 1 (healthy) and Cluster 4 (faltering). The *Klebsiella* finding was also supported by a higher abundance in Cluster 2 compared to Cluster 1. Finally, *Sutterella*, *Peptostreptococcus*, and *Enterococcus* were higher in abundance in Cluster 4 compared to Cluster 1 while *Terrisporobacter* and *Haemophilus* were in higher abundance in Cluster 4 compared to Cluster 3. Additionally, several taxa were identified at 12 months and are presented in the supplementary material (Table S7). All taxa reported were significant after false discovery rate (FDR) correction.

We then pooled all samples to evaluate which taxa identified above were generally positively or negatively correlated with adiposity-related metrics (FMZ, WAZ, WHZ) or physiological/structural-related metrics (FFMZ, HAZ) across different timepoints. First, we evaluated taxa higher in abundance in the growth faltering subgroup at 6 months. We evaluated the correlation between these taxa at each timepoint with physiological/structural-related metrics at equivalent or later timepoints, when relevant. Figure 4A suggests that these taxa at 4 and 6 months were predominately negatively correlated with FFMZ and HAZ at 6 and 12 months. These taxa, many of which are known pathobionts, likely contribute to a pro-inflammatory gut environment that impairs nutrient absorption and diverts host energy from lean mass accretion and linear growth towards managing inflammation. Second, we evaluated taxa higher in abundance in the catchup growth subgroup or overall beneficial taxa (e.g., *Megasphaera*, *Lactobacillus*, *Bifidobacterium*). Given the role of adiposity utilization during both catchup growth and healthy development, we evaluated the correlation between these taxa at each timepoint with adiposity-related metrics at equivalent or later timepoints. Figure 4B suggests that several facultative anaerobes hypothesized to aid in a catchup growth transition were positively correlated at key points with FMZ, WAZ, and WHZ, but this positive association may have been limited to certain timeframes. For example, a higher abundance of *Klebsiella* at 4 and 6 months was positively associated with all three adiposity related metrics at 6 months. However, *Klebsiella* abundance at 6 and 12 months were negatively associated with adiposity metrics at 12 months. Finally, taxa related to healthy growth were predominantly positively correlated with adiposity-related metrics across timepoints.

**Figure 4.**
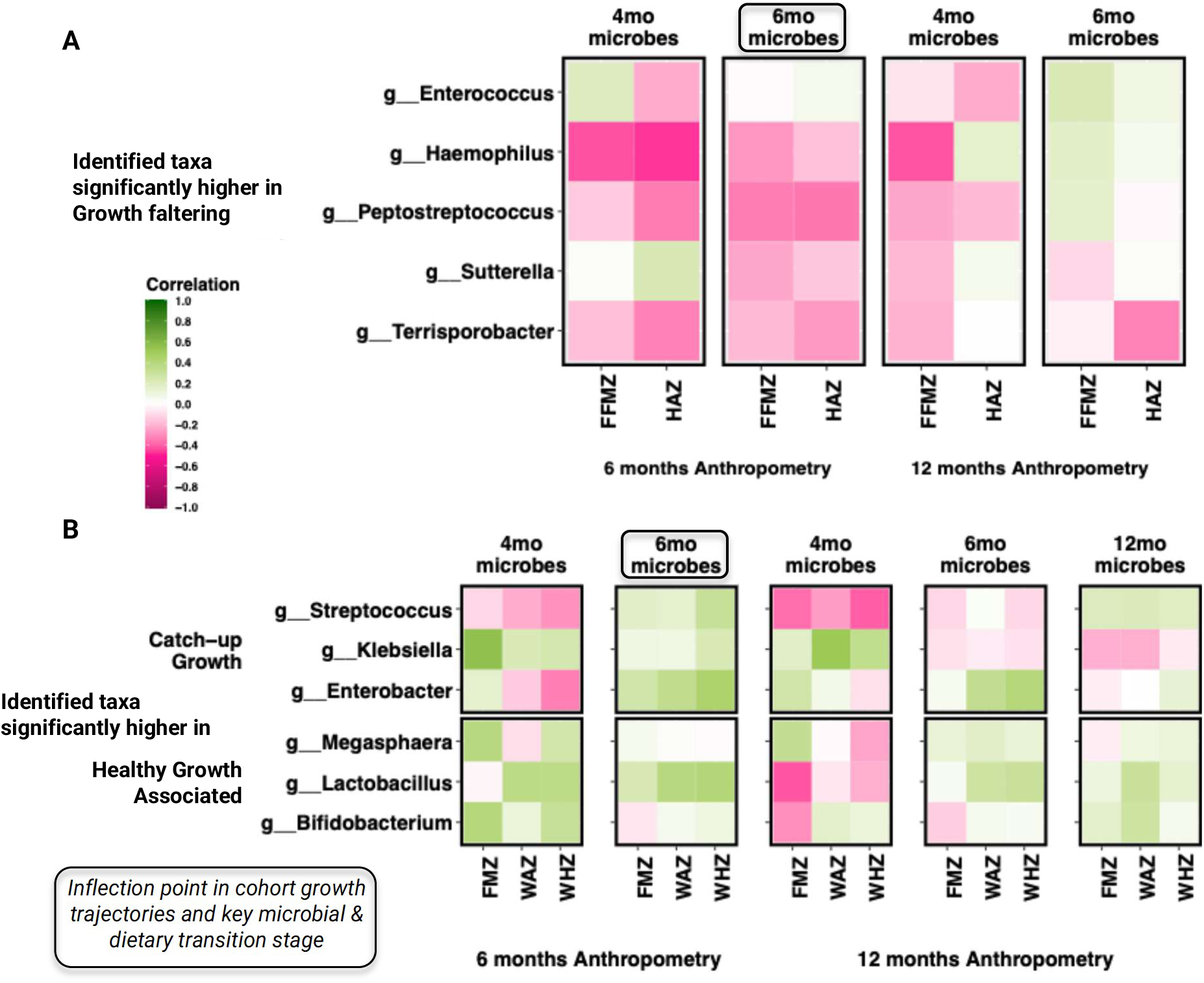
Heatmap of Pearson correlation analysis of relevant taxa (differentially abundant between clusters identified in Table S7) with body composition and anthropometrics at each timepoints. **A)** Growth faltering associated taxa and **B)** taxa associated with catchup growth and healthy growth. The correlations are indicated by colors (green: positive; pink: negative). The color represents the effect size and direction of the correlation. The intensity of the color reflects the correlation coefficient. 4mo = 4months; 6mo= 6months; 12mo=12months. The relationship between 12mo microbes and 12mo FFMZ and HAZ were not evaluated due to the known temporal delay in the relationship.

### Healthy Gut Microbes were Negatively Associated with Morbidity

Finally, we evaluated associations between morbidity across subgroups. Figure 5A depicts the morbidity index across clusters with Cluster 4 having the highest rate. Of the 3 categories that make up the morbidity index (diarrhea, cough/wheezing, runny nose/sneezing), Cluster 4 differences were most pronounced in the cough/wheezing and runny nose/sneezing categories (see Supplementary Figure S12). For water, sanitation, or hygiene indices, there were no discernable patterns or significant differences across clusters (see Supplementary Figure S13).

**Figure 5.**
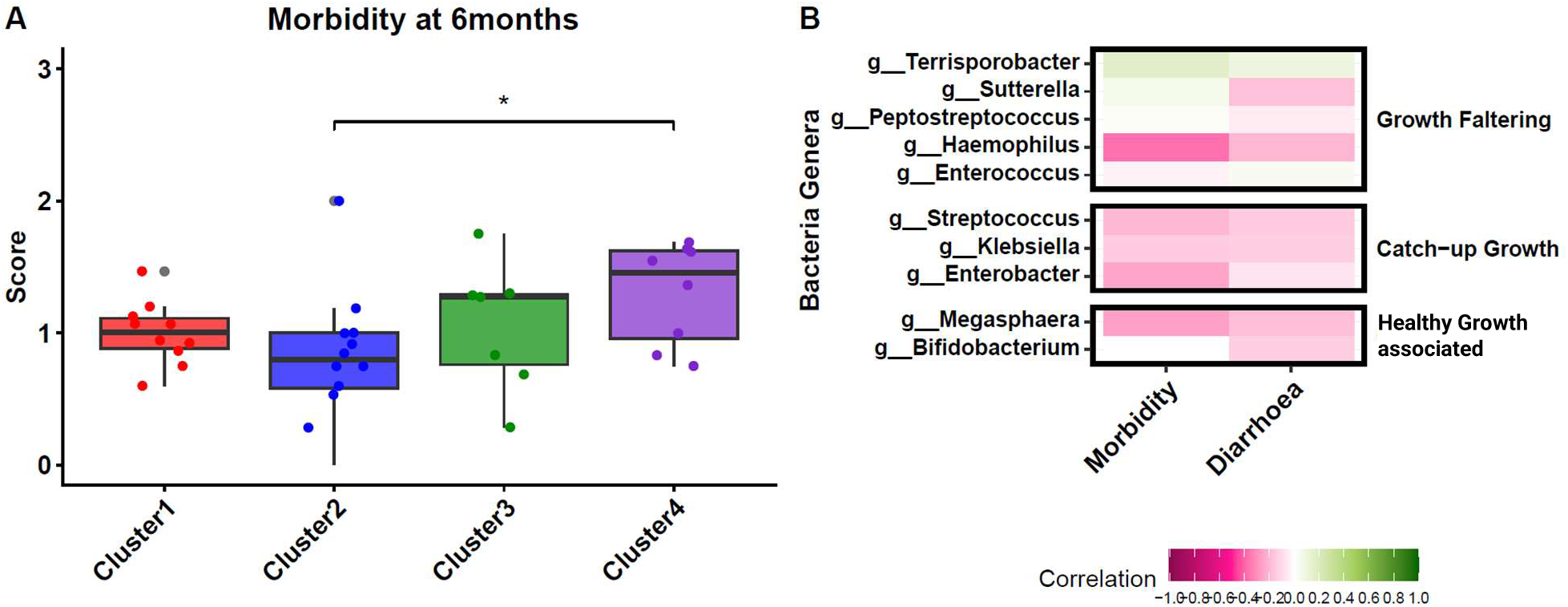
(A) Plot of Morbidity Scores by cluster at 6 months. (B) Heatmap of key taxa that correlates with morbidity and diarrhea at 6 months. The correlations are indicated by colors (green: positive; pink: negative). The color represents the effect size and direction of the correlation. The intensity of the color reflects the correlation coefficient. * p value < 0.05; Wilcoxon rank sum tests were calculated, and all p-values were corrected with Benjamini-Hochberg post-hoc corrections.

We then pooled all samples to evaluate the correlation between the abundance of taxa identified as abundant for each cluster at 6 months and morbidity and diarrhea at 6 months (Figure 5B). All taxa identified that were higher in abundance in catchup and healthy clusters had a negative correlation with morbidity and diarrhea, while in the taxa related to the growth faltering subgroup, had a mix of positive and negative correlations.

## DISCUSSION

Integrating longitudinal growth trajectories with body composition and gut microbiome data from our Soweto cohort revealed four distinct early-life growth phenotypes that remain obscured when growth is evaluated at the cohort level. By clustering children based on parameters from the Jenss-Bayley growth model, we identified phenotypically meaningful subgroups - characterized as healthy growth, stunted but recovering, catch-up growth, and growth faltering - that provide a more nuanced assessment than aggregate anthropometric summaries. These phenotypes diverged primarily within the first 6 months of life. This supports recent evidence that suggests the most significant postnatal growth rate changes occur within the first 3–6 months (Benjamin-Chung et al., 2023), while we extend these findings by linking these divergent paths to unique physiological characteristics. Specifically, infants in Cluster 4 exhibited an interesting divergence between linear growth trajectories and the preservation of core tissues. While their HAZ remained low but consistent between 6 and 12 months, these infants maintained a relatively constant fat-free mass accumulation (FFMZ) despite a simultaneous downward trend in fat mass accumulation (FMZ). This pattern suggests a possible ’triage’ of available energy, where fat stores may be mobilized to support the maintenance of essential organ and muscle mass (FFM) when nutritional or environmental resources are insufficient to support a standard linear growth velocity. The lack of correlation between FFMZ and HAZ across the cohort further supports the hypothesis that a seemingly stable FFMZ in faltering infants may not necessarily indicate robust health. Instead, it may reflect a successful, albeit metabolic costly, adaptation where the body prioritizes the integrity of existing lean tissues over the high energy demands of compensatory bone growth. These findings suggest that in high-burden environments, evaluating body composition alongside linear growth may reveal underlying metabolic trade-offs that are not apparent through anthropometry alone (Momberg et al., 2023).

This potential metabolic triage may also be compounded by a distinct microbial environment that imposes a chronic ’inflammatory tax’ on the developing infant. In the growth-faltering phenotype (Cluster 4), we observed a relative enrichment of opportunistic pathobionts, most notably *Paraclostridium*, *Terrisporobacter*, and *Haemophilus*. The presence of *Paraclostridium*, a genus frequently associated with dysbiotic states and systemic infection (Edagiz et al., 2015; Focardi et al., 2025), may suggest a pro-inflammatory gut environment that further diverts energy away from linear growth toward immune activation. These findings align with the physiological costs observed in Cluster 4, where the metabolic burden of managing potential pathobiont overgrowth may necessitate the mobilization of energy reserves. Conversely, children in the healthy growth group (Cluster 1) appeared to maintain an environment more conducive to nutrient utilization, characterized by an enrichment of functional taxa categorized as “beneficial fermenters,” such as *Megasphaera* and *Parabacteroides*. *Megasphaera* is recognized for its production of short-chain fatty acids (SCFAs) that support metabolic health (Canani, 2011; Fluitman et al., 2017), while *Parabacteroides* has been linked to the degradation of complex carbohydrates and the regulation of gut inflammation (Cui et al., 2022). Furthermore, the depletion of these functional taxa in Cluster 4 may suggest a loss of the protective microbial ’shield’ that usually facilitates healthy nutrient partitioning and, subsequently, linear growth.

The catch-up growth phenotype (Cluster 3) may represent a distinct biological state characterized by a unique microbial succession that facilitates rapid recovery. This signature was marked by a potential transient enrichment of facultative anaerobes, specifically ’opportunistic energy harvesters’ such as *Klebsiella* and *Enterobacter*, which were most prominent around 6 months of age. These taxa may be functionally adaptive, as these organisms are better equipped to thrive in the partially oxygenated or mildly inflammatory gut environments common in high-exposure settings (Senn et al., 2020). By maintaining energy harvest under these sub-optimal conditions, these opportunistic taxa may provide a metabolic bridge that enables the transition from initial faltering toward improved linear growth. However, the presence of such energy-efficient taxa during this critical window underscores the importance of the quality of catch-up growth. While these microbes may unlock the energy required for HAZ recovery, the long-term metabolic health of the infant likely depends on whether this ’bridge’ leads to a balanced accretion of lean mass (FFM) or a disproportionate accumulation of fat mass (FM), the latter of which can increase later-life risks for non-communicable diseases.

The alignment of growth trajectories with body composition, microbiome profiles, and morbidity suggests a model in which early-life growth outcomes emerge from dynamic interactions between microbial ecology, energy allocation, and inflammatory burden. Our findings are consistent with a gut-growth feedback loop wherein infants experiencing growth-faltering (Cluster 4) face a high pathobiont burden and elevated morbidity rates, particularly respiratory symptoms, which may induce a chronic pro-inflammatory state. This environment appears to necessitate a ’metabolic triage’ strategy: the body seemingly prioritizes the maintenance of essential fat-free mass over the high energy demands required to maintain fat-mass accretion. This pattern aligns with broader life history frameworks of developmental energy partitioning, in which fat mass during infancy functions less as a passive marker of energy deficit and more as a metabolically flexible reserve that can be adaptively mobilized to sustain vital tissues and high-priority organ development under energetic constraint (Said-Mohamed et al. 2018). In addition, the transient enrichment of ’opportunistic energy harvesters’ in the catch-up group (Cluster 3) suggests an adaptive, transitional microbial state that may provide the metabolic bridge required for recovery under similarly constrained conditions. Ultimately, these divergent paths are defined not just by calorie intake, but by the physiological and microbial capacity to manage the metabolic costs of a high-exposure environment, setting the foundational trajectory for long-term health and development.

While this study provides a high-resolution, multi-dimensional view of infant development, it is limited by a relatively small sample size (n=45) and the use of 16S rRNA amplicon sequencing. These constraints restrict our ability to identify specific microbial strains or directly evaluate functional gene pathways that might further explain the metabolic differences between phenotypes. Despite these limitations, our trajectory-based framework offers a biologically grounded approach for identifying the mechanisms that drive growth faltering and recovery. Our findings suggest that future “microbiota-sensitive” interventions should move beyond population-level summaries to target these specific states. For instance, children stuck in a cycle of metabolic triage may benefit from interventions that reduce pathobiont-induced inflammation or supplement “beneficial fermenters” like *Megasphaera* to restore SCFA production and support lean mass accretion.

## CONCLUSIONS

This study demonstrated that integrating longitudinal data on growth, body composition, and the gut microbiome can uncover distinct phenotypes of child growth that are not apparent through anthropometry alone. By identifying key microbial taxa and diversity patterns associated with healthy, catch-up, and faltering growth, we provide new insights into the potential microbial contributors to, or consequences of, early-life growth trajectories. The presence of facultative anaerobes during catch-up growth suggests a transitional microbial state that may reflect physiological recovery under partial inflammatory conditions. Importantly, associations between microbial taxa and FFMZ or linear growth indicators emphasize the need to consider growth as a multifaceted biological process. These findings support the value of microbiome-informed approaches for stratifying risk and guiding targeted interventions to promote healthy growth in early life.

## Data Availability Statement

Data is available upon reasonable request from the authors.

## Acknowledgment

The authors are grateful for the communities and field workers for which this work would not have been possible.

