## Supplemental Material for "From Growth Faltering to Recovery: Gut Microbial and Body Composition Signatures of Early Childhood Malnutrition Phenotypes"

### Supplementary Figures

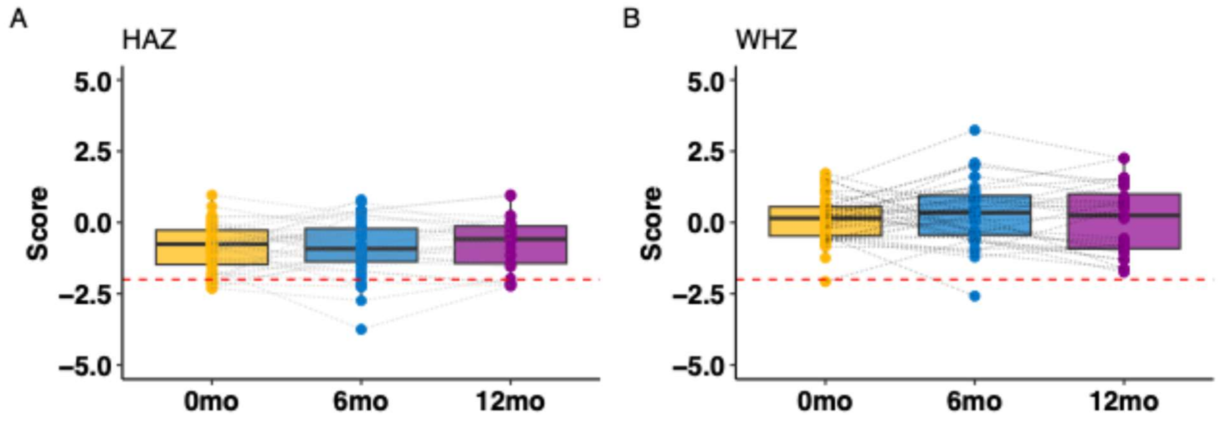

**Figure S1**. HAZ and WHZ measurements at birth (0month), 6 months, and 12 months. HAZ and WHZ did not change significantly from birth to 12 months. 0mo = 0months; 6mo= 6months; 12mo=12months.

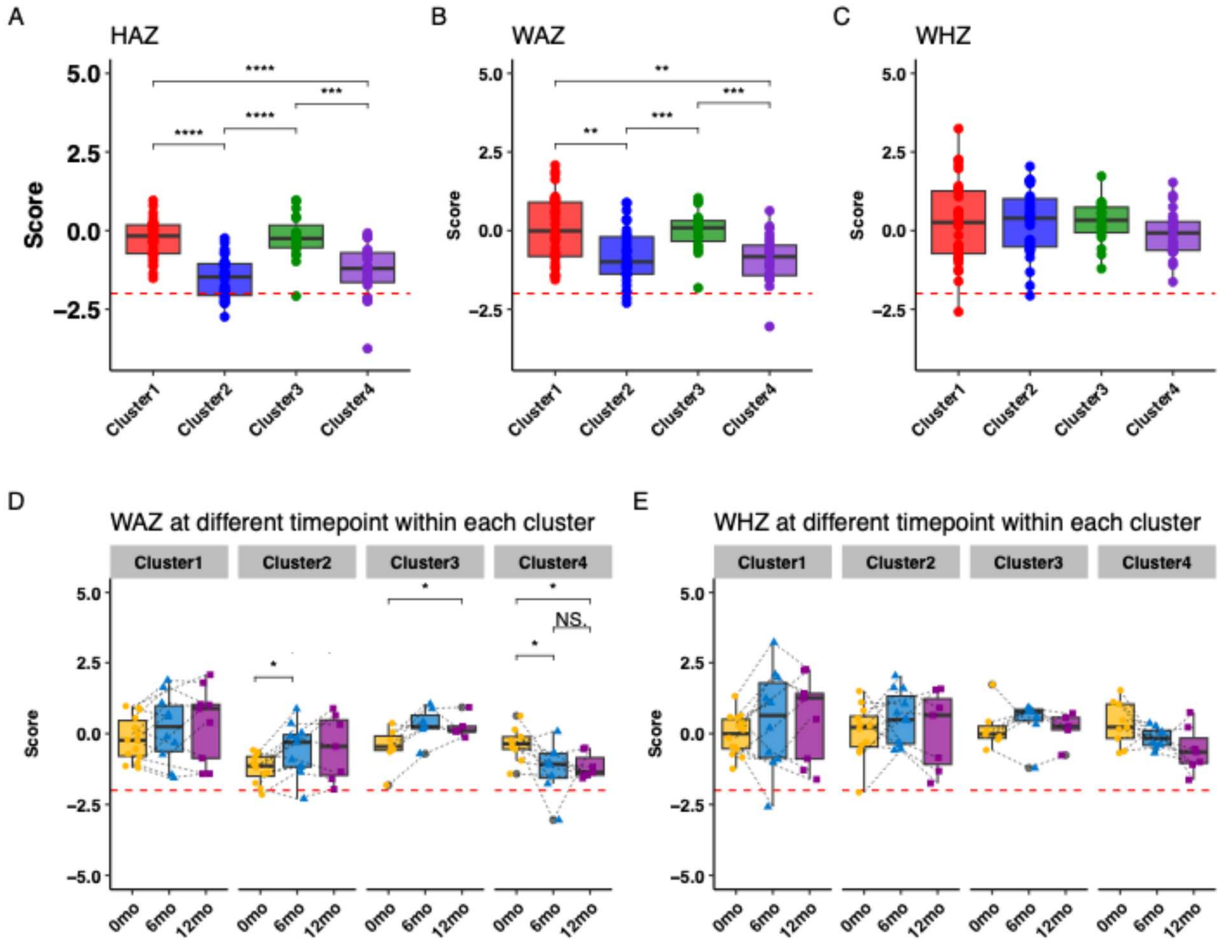

**Figure S2.** Comparison of the three growth forms between clusters and timepoints. A) HAZ, B) WHZ, C) WAZ, D) WAZ by timepoint, E) WHZ by timepoint.  * p value < 0.05; ** p value < 0.01; **** p value < 0.0001; Wilcoxon rank sum tests were calculated, and all p-values were corrected with Benjamini-Hochberg post-hoc corrections.

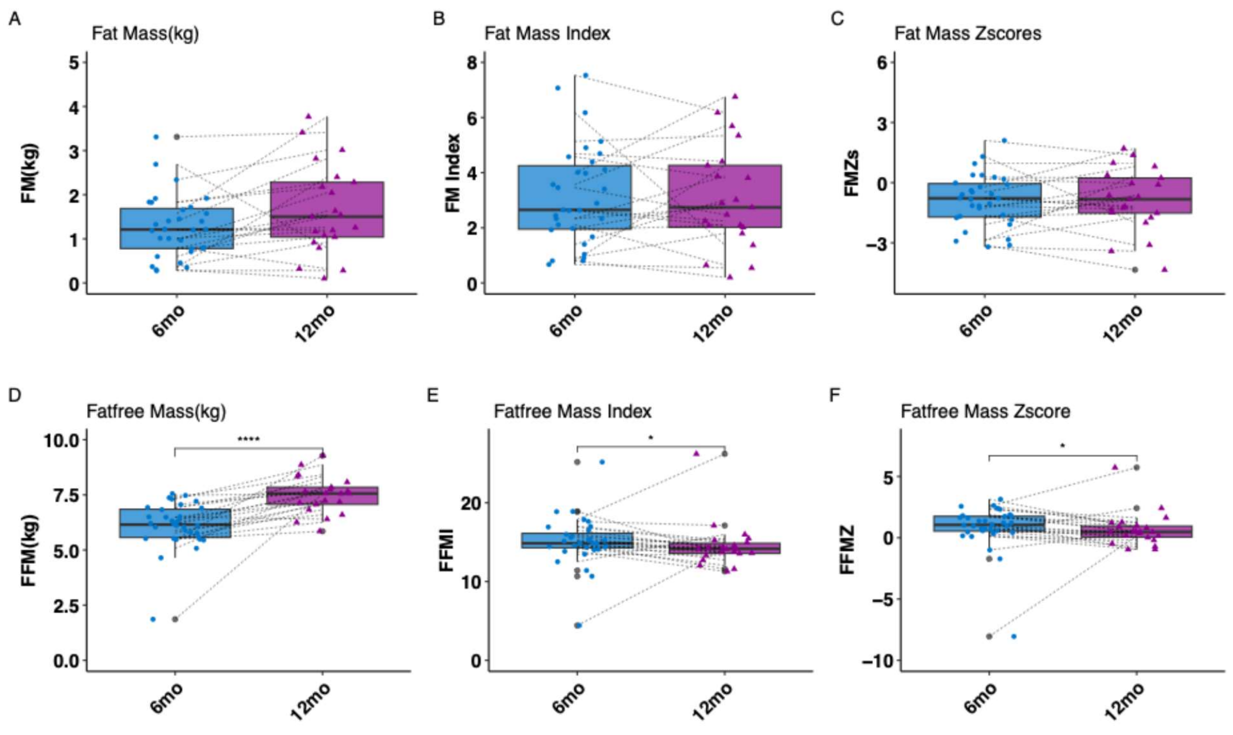

**Figure S3**. Overall Body composition plot by Timepoint. A) Fatmass B) Fatmass index C) Fatmass zscore D) Fatfree mass E) Fatfree Mass Index F) Fatfree mass zscore. 6mo= 6months; 12mo=12months. **** p value < 0.0001; * p value < 0.05; t.tests were calculated, and all p-values were corrected with Benjamini-Hochberg post-hoc corrections.

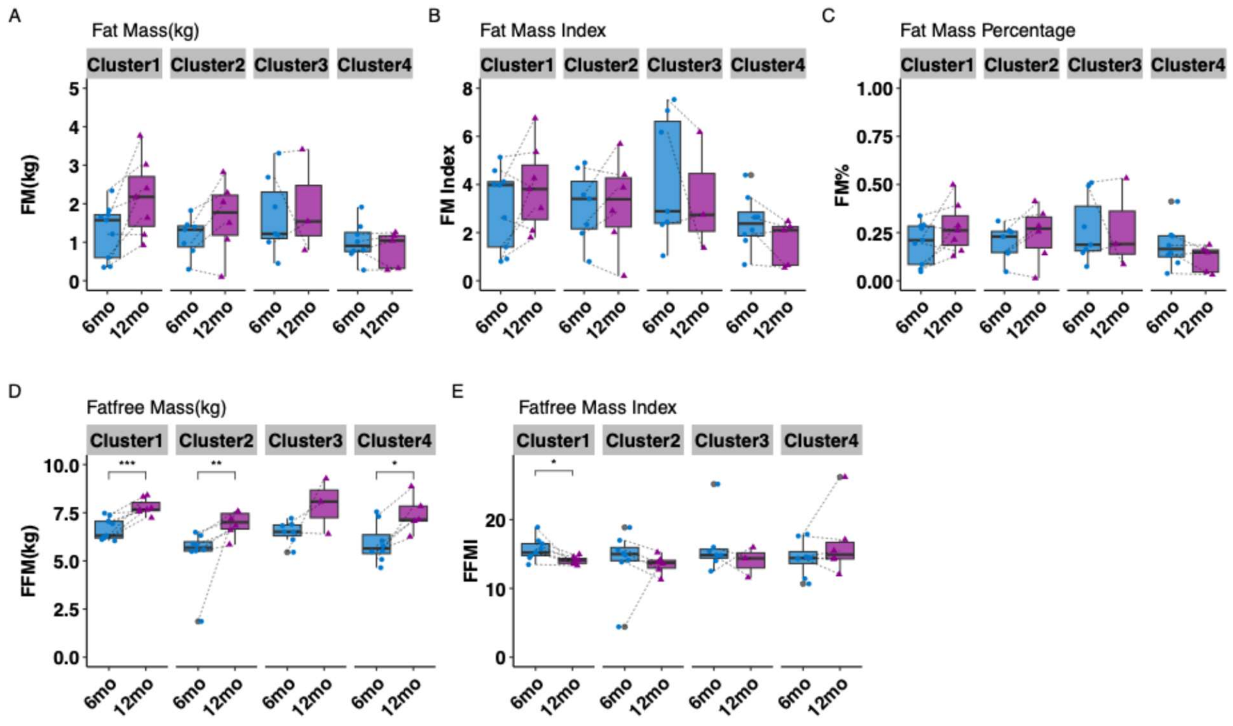

**Figure S4**. Body composition plot by Timepoint. A) Fatmass B) Fatmass index C) Fatmass percentage D) Fatfree mass E) Fatfree Mass Index. 6mo= 6months; 12mo=12months. *** p value < 0.001; * p value < 0.05; t.tests were calculated, and all p-values were corrected with Benjamini-Hochberg post-hoc corrections.

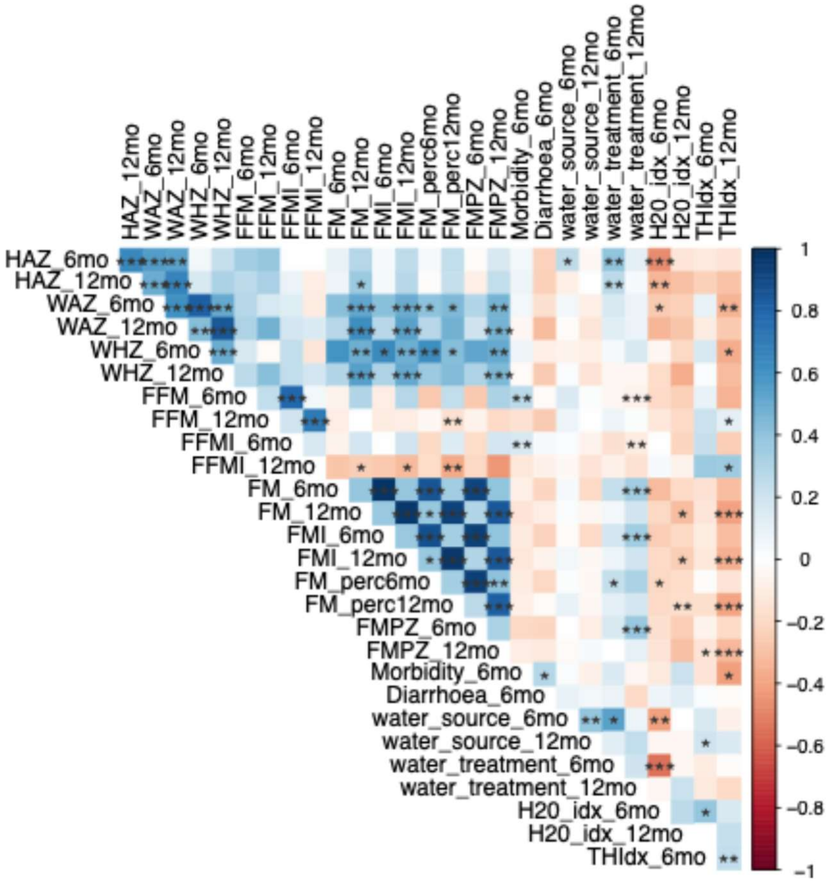

**Figure S5.** Heatmap of spearman correlation analysis among WASH, anthropometric, body composition and morbidity. The correlations are indicated by colors (blue: positive; red: negative). The color represents the effect size and direction of the correlation. The intensity of the color reflects the correlation coefficient. Significant correlations are indicated by *p<0.05, **p<0.01, ***p<0.001

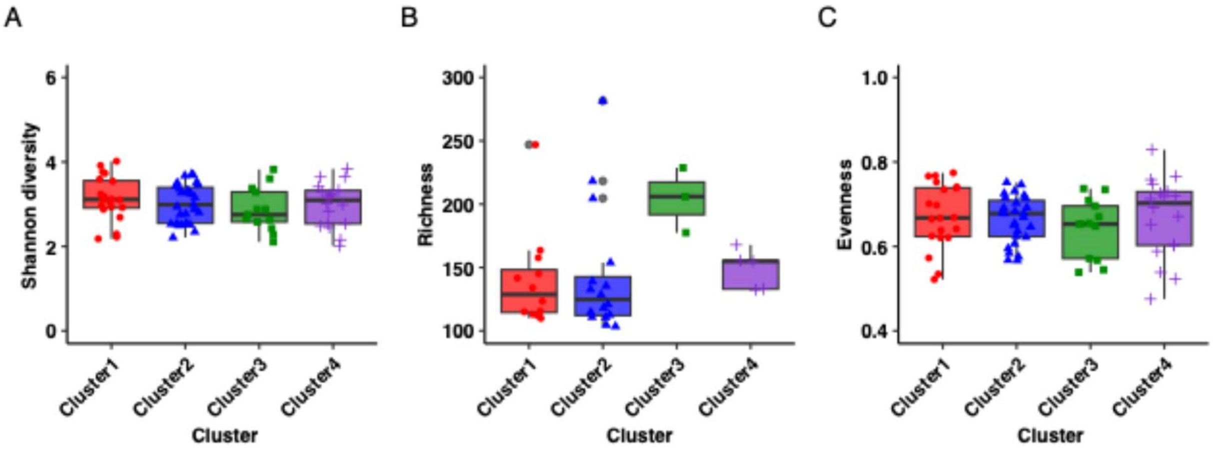

**Figure S6.** Diversity comparisons between clusters A) richness , B) Shannon index, and C) evenness and timepoints D) richness , E) Shannon index, and F) evenness across microbiome composition. Wilcoxon rank sum tests were performed for all tests with Benjamini-Hochberg post-hoc corrections for multiple comparisons. 0mo = 0month; 6mo = 6months; 12mo = 12months.

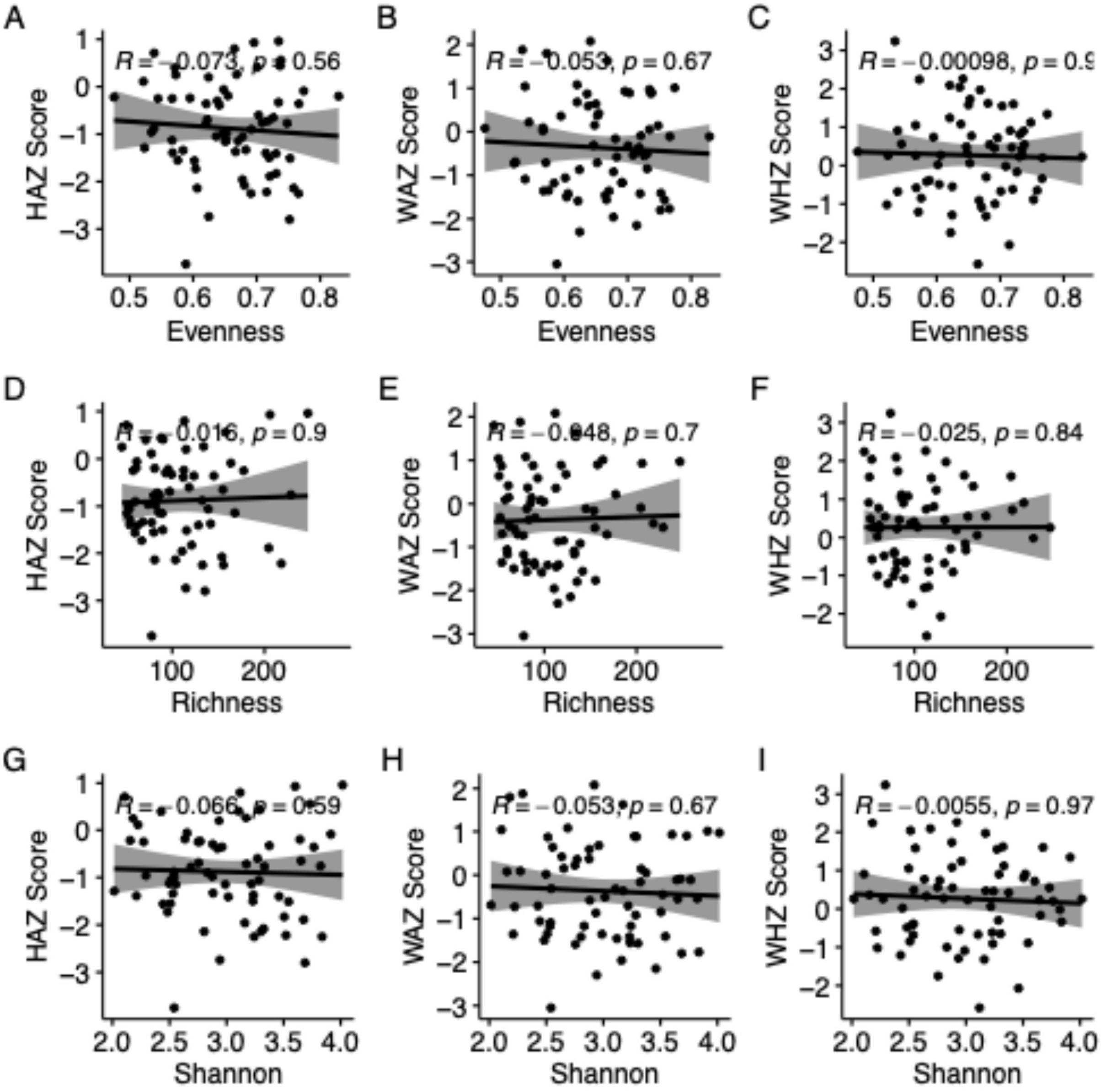

**Figure S7.** Correlation analysis between alpha diversity and anthropometrics. R value represents the strength and direction of the correlation. P values indicates statistical significance.

A

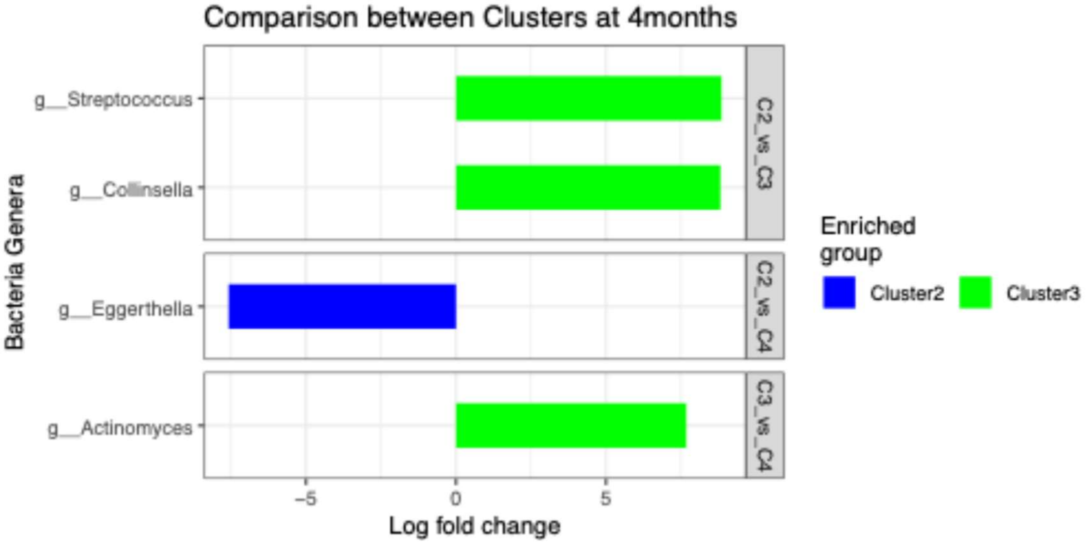
Comparison between Clusters at 4months
g__Streptococcus
g__Collinsella
g__Eggerthella
g__Actinomyces
C2_vs_C3
C2_vs_C4
C3_vs_C4
Bacteria Genera
−5
0
5
Log fold change
Enriched group
Cluster2
Cluster3

B

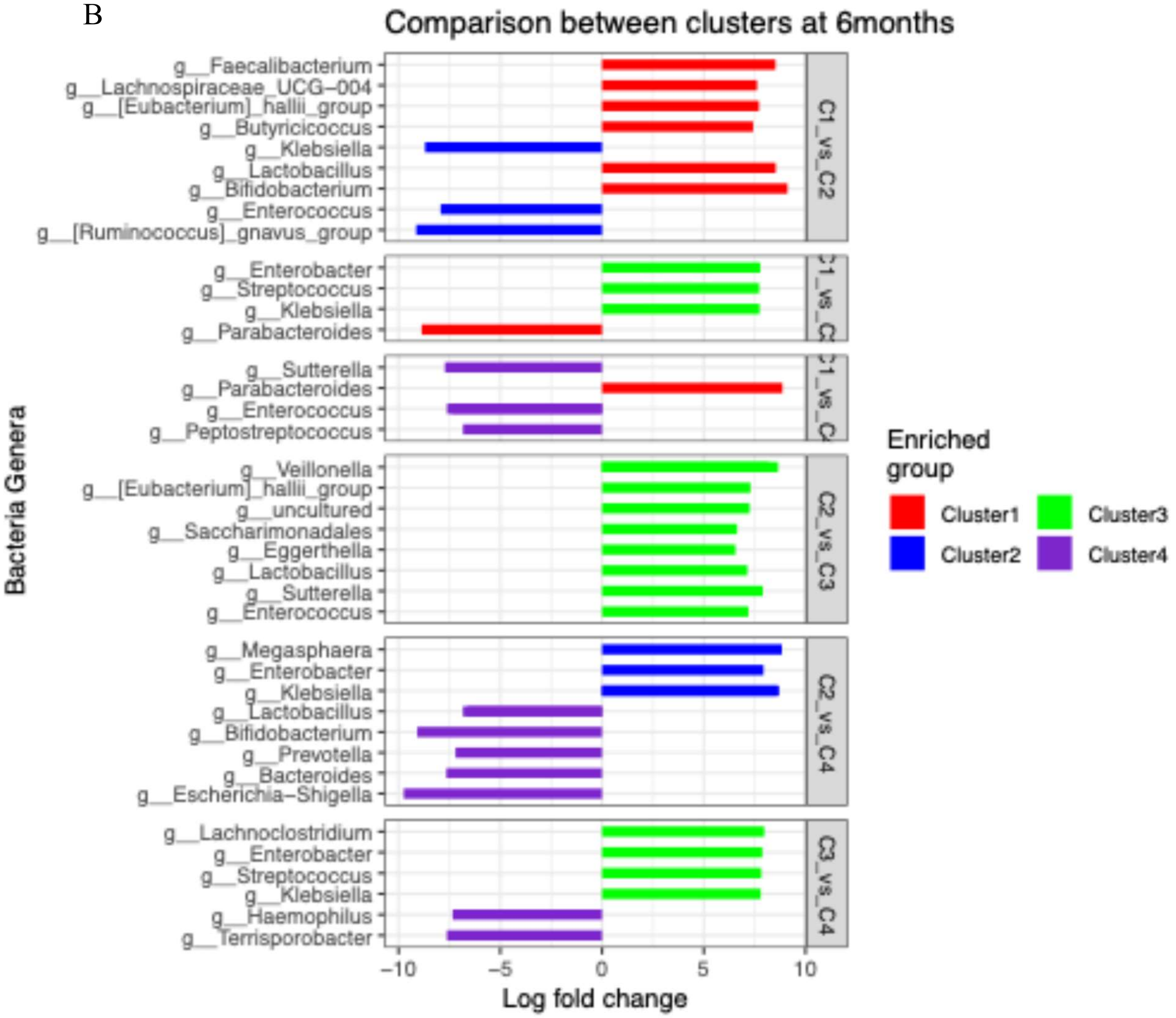
Comparison between clusters at 6months
g__Faecalibacterium
g__Lachnospiraceae_UCG−004
g__[Eubacterium]_hallii_group
g__Butyricicoccus
g__Klebsiella
g__Lactobacillus
g__Bifidobacterium
g__Enterococcus
g__[Ruminococcus]_gnavus_group
g__Enterobacter
g__Streptococcus
g__Klebsiella
g__Parabacteroides
g__Sutterella
g__Parabacteroides
g__Enterococcus
g__Peptostreptococcus
g__Veillonella
g__[Eubacterium]_hallii_group
g__uncultured
g__Saccharimonadales
g__Eggerthella
g__Lactobacillus
g__Sutterella
g__Enterococcus
g__Megasphaera
g__Enterobacter
g__Klebsiella
g__Lactobacillus
g__Bifidobacterium
g__Prevotella
g__Bacteroides
g__Escherichia−Shigella
g__Lachnoclostridium
g__Enterobacter
g__Streptococcus
g__Klebsiella
g__Haemophilus
g__Terrisporobacter
C1_vs_C2
C2_vs_C3
C2_vs_C4
C3_vs_C4
Bacteria Genera
−10
−5
0
5
10
Log fold change
Enriched group
Cluster1
Cluster2
Cluster3
Cluster4

C

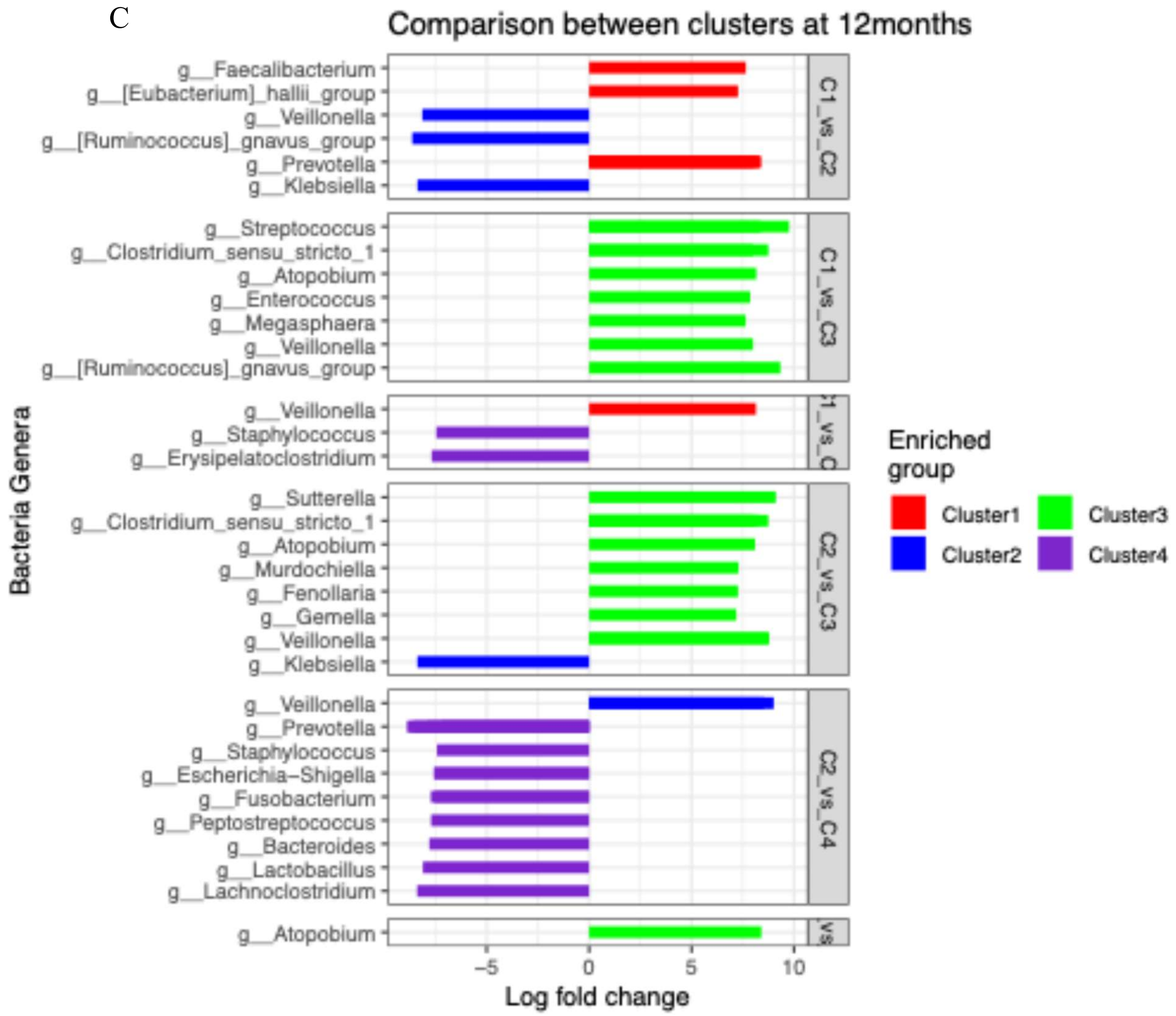

**Figure S8.** Microbes differentially abundant between cluster groups within timepoint. A) 4months timepoint B) 6months timepoint and C) 12months timepoint. The x-axis shows the log fold change representing the degree of differences in the relative abundance of the microbes between the cluster groups expressed in logarithmic scale. Only taxa having a $p<0.05$ and log fold change >1 are shown. The y-axis shows differentially abundant microbes between the clusters and is colored based on the cluster in which they are most enriched. C1 = Cluster 1; C2= Cluster 2; C3= Cluster 3; C4=Cluster 4.

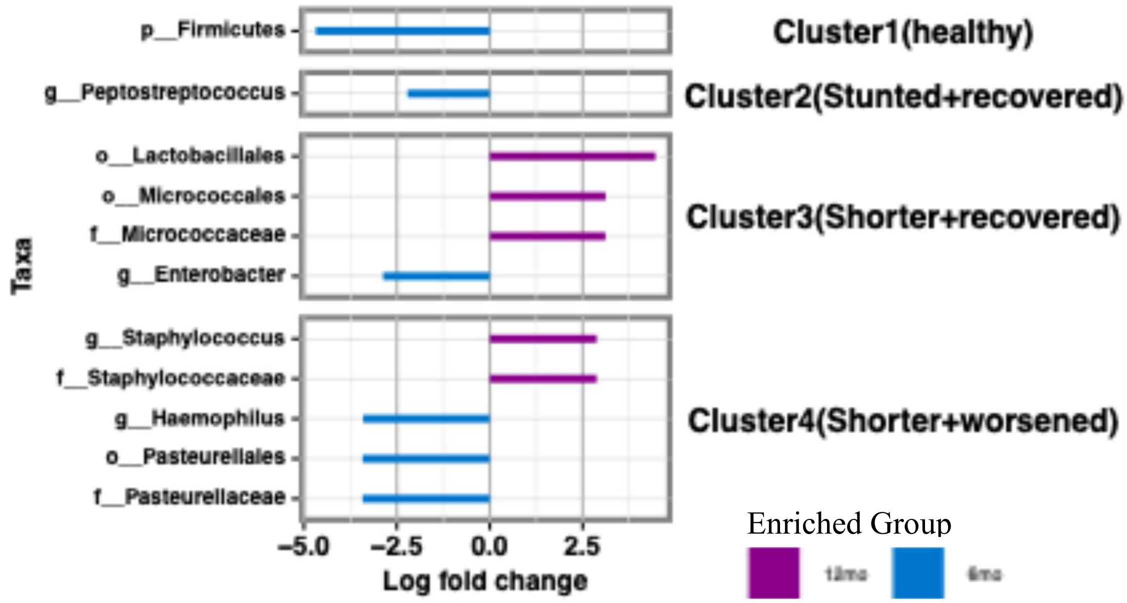

**Figure S9.** Microbes differentially abundant between timepoints within cluster groups. The x-axis shows the log fold change representing the degree of differences in the relative abundance of the microbes between the cluster groups expressed in logarithmic scale. Only taxa having a $p<0.05$ and log fold change >1 are shown. The y-axis shows differentially abundant microbes between the timepoints within the cluster groups and is colored based on the timepoint in which they are most enriched. 6mo = 6month; 12mo = 12months.

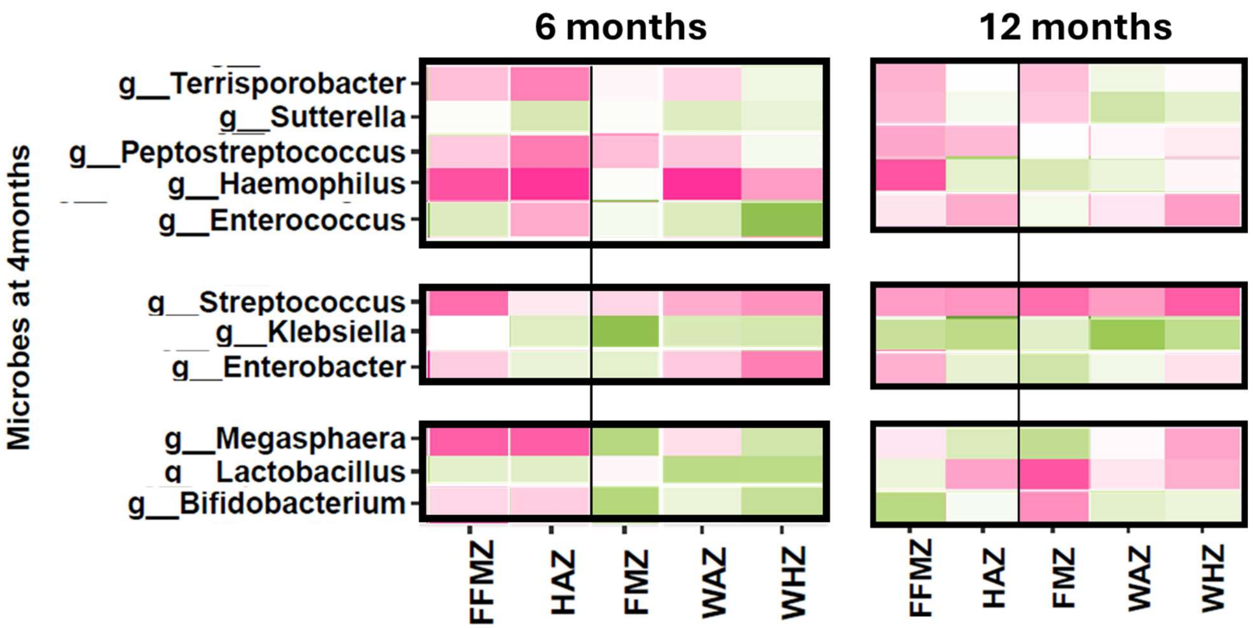

**Figure S10**. Heatmap of specific taxa in higher abundance within specific clusters and their association with body composition and anthropometry. Taxa data are from the 4 month time point and body composition/anthropometry are from the 6 and 12 month time points.

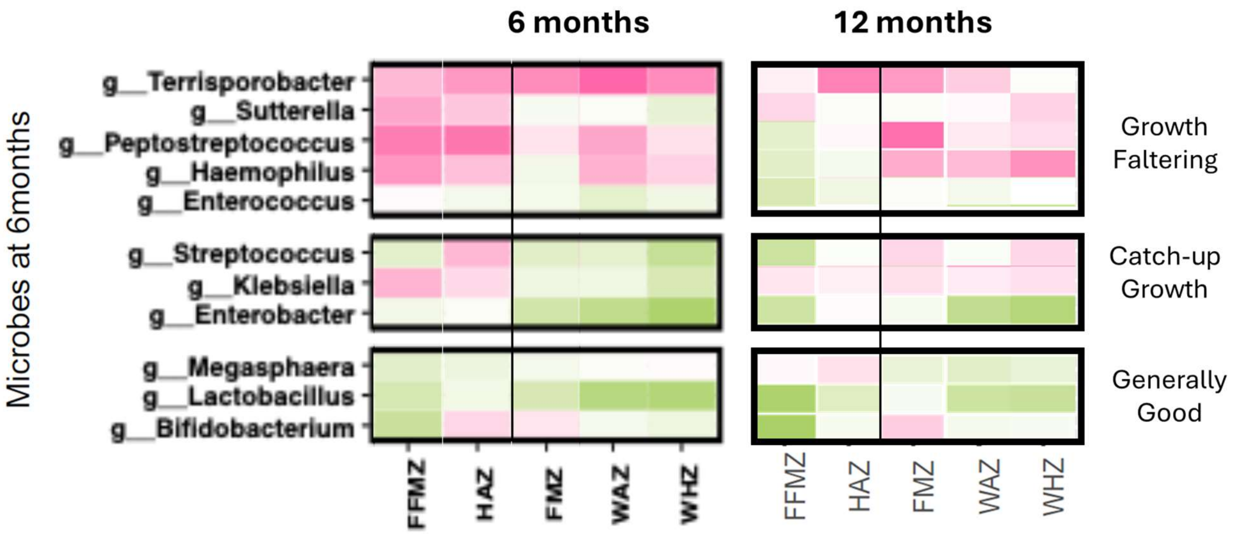

**Figure S11**. Heatmap of specific taxa in higher abundance within specific clusters and their association with body composition and anthropometry. Taxa data are from the 6 month time point and body composition/anthropometry are from the 6 and 12 month time points.

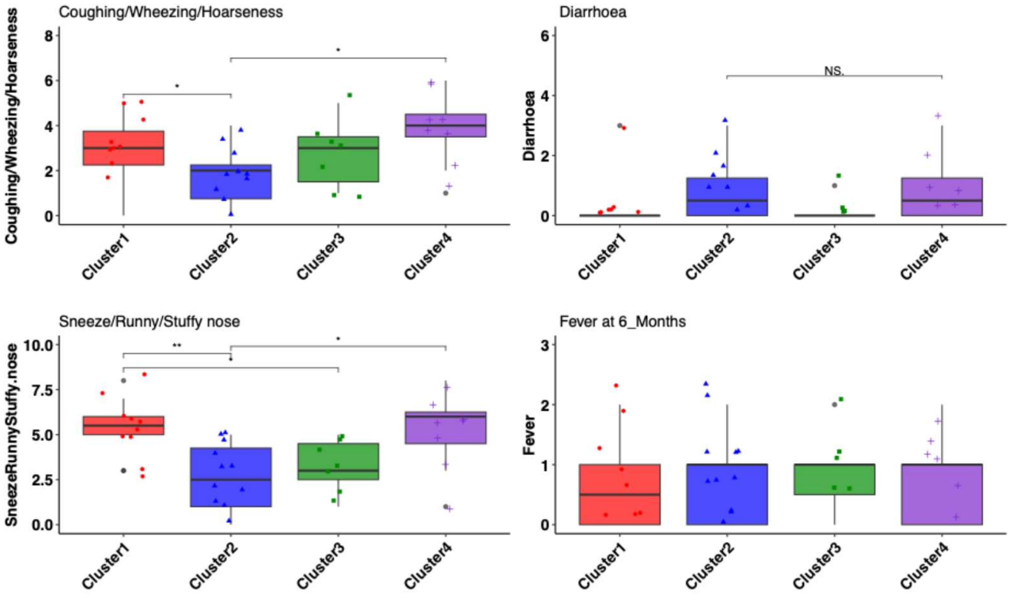

**Figure S12**. A breakdown of the 4 categories that make up with morbidity index, broken out by cluster.

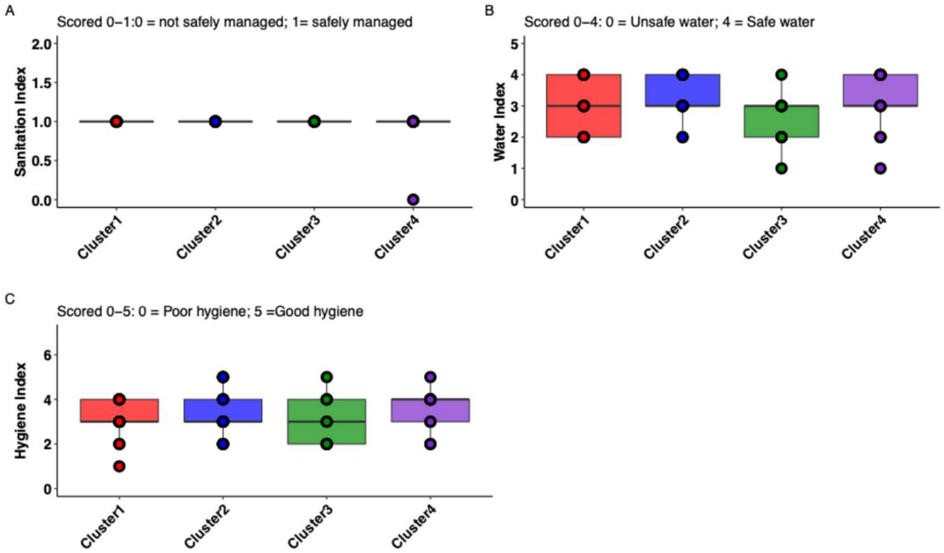

**Figure S13**. Water, sanitation, and hygiene indices for each cluster.

### Supplementary Table

**Table S1.** Characteristics of infants at birth in the Soweto Baby WASH study. ± values indicate standard deviation(sd) from the mean.

| | |
|---|---|
| No. of infants | 45 |
| Sex | Male: 20<br>Female:25 |
| HAZ | 0mo:-0.93±0.8; 6mo:-0.94±0.8; 12mo: -0.91±0.8 |
| WAZ | 0mo:-0.55±0.8; 6mo:-0.55±0.8; 12mo:-0.49±0.8 |
| WHZ | 0mo:0.19±0.8, 6mo:0.20±0.7; 12mo:0.26±0.7 |

**Table S2.** Summary of the average values of anthropometric data in each cluster group at 0, 6, and 12months. ± values indicate standard deviation(sd) from the mean.

| **Clusters** | **HAZ** | | | **WAZ** | | | **WHZ** | | |
|---|---|---|---|---|---|---|---|---|---|
| | 0mo | 6mo | 12mo | 0mo | 6mo | 12mo | 0mo | 6mo | 12mo |
| 1 | -0.25±0.6 | -0.26±0.7 | -0.27±0.5 | -0.18±0.8 | 0.168±1.2 | 0.39±1.3 | -0.04±0.7 | 0.46±1.8 | 0.58±1.5 |
| 2 | -1.82±0.6 | -1.47±0.7 | -1.22±0.7 | -1.27±0.5 | -0.57±0.9 | -0.50±1.2 | 0.06±0.9 | -0.56± 0.9 | 0.11±1.4 |
| 3 | -0.82±0.6 | 0.07±0.5 | 0.24±0.6 | -0.47±0.7 | 0.32±0.6 | 0.25±0.4 | 0.19±0.7 | 0.44±0.8 | 0.18±0.5 |
| 4 | -0.77±0.6 | -1.60±1 | -1.51±0.6 | -0.37±0.6 | -1.22±0.9 | -1.15±0.5 | 0.34±0.8 | -0.15±0.4 | -0.56±0.8 |

**Table S3.** Summary of the average values of WASH factors in each cluster group at 6 and 12months. ± values indicate standard deviation(sd) from the mean.

| Clusters | **Total hygiene** | | **Water index** | | **Sanitation index** | |
|---|---|---|---|---|---|---|
| | **6mo** | **12mo** | **6mo** | **12mo** | **6mo** | **12mo** |
| 1 | 3.25±0.7 | 2.86±1.1 | 3±0.9 | 3±0.7 | 1 | 1 |
| 2 | 3.4±1 | 3.3 | 3.4±0.8 | 3 | 1 | 1 |
| 3 | 2.6±1.4 | 3±01.4 | 2.4±0.9 | 2.8±00.4 | 1 | 1 |
| 4 | 3.6±1.1 | 3.5±0.8 | 3±1 | 3.4±0.8 | 0.9±0.3 | 1 |

**Table S4.** Summary of the average values of morbidity index and illness in each cluster group at 6months. ± values indicate standard deviation(sd) from the mean.

| **Clusters** | **Morbidity** | **Diarrhea** | **Fever** |
|---|---|---|---|
| 1 | 1±0.2 | 0.3±0.9 | 0.7±0.8 |
| 2 | 0.82±0.5 | 0.83±1.0 | 0.83±0.7 |
| 3 | 0.94±0.4 | 0.17±0.4 | 0.83±0.7 |
| 4 | 1.3±0.3 | 0.87±1.1 | 0.75±0.7 |

**Table S5.** Summary of the average values of body composition in each cluster group at 6 and 12months. ± values indicate standard deviation(sd) from the mean.

| Clusters | **FM(kg)** | | **FM Index** | | **FM zscore** | | **FM Percentage** | |
|---|---|---|---|---|---|---|---|---|
| | 6mo | 12mo | **6mo** | **12mo** | **6mo** | **12mo** | **6mo** | 12mo |
| 1 | 1.29±0.7 | 2.23±1.2 | 3.06±0.9 | 3±0.7 | -0.86±1.4 | -0.23±1.6 | 0.19±0.1 | 0.28±0.1 |
| 2 | 1.55±1.5 | 1.56±1.0 | 3.92±3.6 | 3 | -0.43±2.0 | -1.12±2.0 | 0.20±0 | 0.23±0.2 |
| 3 | 1.79±1.1 | | 2.4±0.9 | 2.8±00.4 | -0.1±1.6 | | 0.28±0.1 | |
| 4 | 1.01±0.5 | 0.48±0.3 | 3±1 | 3.4±0.8 | -1.31±1.0 | -2.86±0.8 | 0.18±0.1 | 0.1±0.0 |
| | **FFM(kg)** | | **FFM Index** | | **FFM zscore** | | | |
| | 6mo | 12mo | **6mo** | **12mo** | **6mo** | **12mo** | | |
| 1 | 6.63±0.6 | 7.91±0.6 | 15.7±1.6 | 13.40±2.3 | 1.74±0.83 | 0.62±0.5 | | |
| 2 | 5.36±1.3 | 6.88±0.7 | 13.79±4.1 | 13.47±1.3 | -0.31±3.1 | -0.19±0.6 | | |
| 3 | 6.43±0.6 | | 16.1±4.2 | 13.98±2.2 | 1.25±0.5 | | | |
| 4 | 5.92±1.0 | 9.97±3.1 | 14.4±2.5 | 16.07±4.7 | 0.54±1.5 | 2.50±2.6 | | |

**Table S6.** Permutational Multivariate Analysis of Variance for microbes using distance matrices (*adonis2*) for Bray-Curtis dissimilarity (*k*=2) between clusters

| **Comparisons** | **$R^2$** | **F** | **pvalue** |
|---|---|---|---|
| Cluster4 vs Cluster3 | 0.05 | 1.57 | 0.01 |
| Cluster4 vs Cluster2 | 0.04 | 1.90 | 0.001 |
| Cluster3 vs Cluster2 | 0.035 | 1.26 | 0.009 |
| Cluster4_vs_Cluster1 | 0.045 | 1.42 | 0.006 |

**Table S7**. Differentially abundant taxa at 4, 6, and 12 months between Cluster 1 (Healthy), 3 (Catch-up), and 4 (Malnourished or Growth Faltering). Taxa derived from Figure S8. Each row shows a bivariate comparison. The column shows the name of the taxa that was in significantly higher abundance for that cluster. The last column summarizes trends identified for each timepoint.

| | **Cluster 1: Healthy** | **Cluster 3: Catch-up** | **Cluster 4: Malnourished** | **Trends** |
|---|---|---|---|---|
| **4mo** | | | | |
| Healthy vs Catch-up | None | None | - | **Healthy**: none<br>**Catch-up**: Actinomyces<br>**Malnourished**: none |
| Healthy vs Malnourished | None | - | None | |
| Catch-up vs Malnourished | - | Actinomyces | None | |
| **6mo** | | | | |
| Healthy vs Catch-up | Parabacteroides | Streptococcus<br>Klebsiella<br>Enterobacter | - | **Healthy**:<br>Parabacteroides<br>**Catch-up**:<br>Streptococcus,<br>Klebsiella, Enterobacter<br>**Malnourished**:<br>Sutterella,<br>Peptostreptococcus, |
| Healthy vs Malnourished | Parabacteroides | - | Sutterella<br>Peptostreptococcus<br>Enterococcus | |
| Catch-up vs Malnourished | - | Streptococcus<br>Klebsiella | Terrisporobacter<br>Haemophilus | |

| | | | | |
|---|---|---|---|---|
| | | Enterobacter<br>Lachnoclostridium | | Enterococcus,<br>Terrisporobacter,<br>Haemophilus |
| **12mo** | | | | |
| Healthy vs Catch-up | None | Streptococcus<br>Veillonella<br>Enterococcus<br>Megasphaera<br>Clostridium-sensu<br>Ruminococcus<br>Atopobium | - | **Healthy**: none<br>**Catch-up**: Atopobium, (Strep remains high vs healthy at 6mo)<br>**Malnourished**: (Veillonella, lack of), Staphylococcus, Erysipolatoclostridium |
| Healthy vs Malnourished | Veillonella | - | Staphylococcus<br>Erysipolatoclostridium | |
| Catch-up vs Malnourished | - | Atopobium<br>Veillonella | None | |
